# Continuous, long-term crawling behavior characterized by a robotic transport system

**DOI:** 10.1101/2023.02.27.530235

**Authors:** James Yu, Stephanie Dancausse, Maria Paz, Tolu Faderin, Melissa Gaviria, Joseph Shomar, Dave Zucker, Vivek Venkatachalam, Mason Klein

## Abstract

Detailed descriptions of behavior provide critical insight into the structure and function of nervous systems. In *Drosophila* larvae and many other systems, short behavioral experiments have been successful in characterizing rapid responses to a range of stimuli at the population level. However, the lack of long-term continuous observation makes it difficult to dissect comprehensive behavioral dynamics of individual animals and how behavior (and therefore the nervous system) develops over time. To allow for long-term continuous observations in individual fly larvae, we have engineered a robotic instrument that automatically tracks and transports larvae throughout an arena. The flexibility and reliability of its design enables controlled stimulus delivery and continuous measurement over developmental time scales, yielding an unprecedented level of detailed locomotion data. We utilize the new system’s capabilities to perform continuous observation of exploratory behavior over a duration of six hours with and without a thermal gradient present, and in a single larva for over 30 hours. Long-term free-roaming behavior and analogous short-term experiments show similar dynamics that take place at the beginning of each experiment. Finally, characterization of larval thermotaxis in individuals reveals a bimodal distribution in navigation efficiency, identifying distinct phenotypes that are obfuscated when only analyzing population averages.

## Introduction

A complete description of an organism’s behavior, or any responsive system more generally, would include a map of how inputs transform into outputs. Reflexes or decisions made by the peripheral and central nervous systems, for example, can be characterized as functions that take surrounding environmental (and internal) stimulus information, process it, and lead to a physical behavior. An organism’s transformation properties are rarely constant, and instead change over short and long time scales, determined by its stimulus history and development. A comprehensive understanding of animal behavior and its underlying mechanisms would ideally address all time scales between fast neural responses through the slow physiological changes associated with development (1–4). Short-term responses to individual stimuli have been characterized in many organisms, but a high-bandwidth treatment with continuous measurement of behavior through the entire time course of an animal’s development has been prohibitive (5–7). While shorter experiments have been successful in characterizing behavior and acute responses to stimuli at the population level, the lack of continuous observation make it difficult to dissect individual behavioral dynamics and their development over time. In most organisms, behavior is too fast, too complicated, is performed in too large of a space, or otherwise too difficult to observe with sufficient resolution over a long time. An ideal system would perform slow, well-defined actions in a confined space while responding robustly to internal and external stimuli.

The *Drosophila melanogaster* larva model system presents an opportunity to study navigational behavioral dynamics over long time scales, well-suited to a detailed quantitative treatment. It possesses a well-defined, slow behavioral repertoire and robust response to many stimuli (8–10). The fruit fly has a relatively short life cycle with high accessibility to their three short larval stages (11). During these stages, larvae are highly food-motivated and thus in the absence of food engage in nearly continuous exploratory movement, which facilitates behavioral studies of locomotion over long times (12). They also demonstrate responses to chemosensory cues (13–15), mechanical and nociceptive stimulation (16–18), light (19), as well as the ability to retain memory and change their behaviors in accordance to learned experiences, and habituate to prolonged stimuli (20– 23). Larvae also perform robust behavioral responses to temperature changes, engaging in thermotaxis along thermal gradients, and modulating aspects of their locomotive behavior, in particular their turning rate, in order to reach optimal conditions (6, 24). The recent availability of a connectome brain wiring diagram (25, 26) and numerous genetic tools have facilitated probing the neural circuits and molecular mechanisms that underlie these behaviors (2, 8, 27–30). Here we focus on directly observed exploration and navigation behavior and seek to continuously measure fly larva crawling over times scales of many hours.

Responses to a wide range of stimuli in larvae typically occur through changes in their navigation and locomotion. Their navigation behavior, when confined to flat 2D spaces, is akin to an organism-scale 2D random walk (31, 32), characterized as an alternating sequence of forward crawling “runs” and direction-altering “turns”, making the animal’s behavior and response to stimuli straightforward to quantify (7, 33–35). However, larvae crawl away and typically remain at the edges of confining barriers, or climb walls or bury into a substrates (9). Either scenario results in the termination of exploratory behavior, limiting most experiments to shorter snap-shots of larval behavior, typically on the order of 10 minutes (6, 7, 36, 37). Manually prolonging exploratory crawling behavior, such as picking up a larva with a wet paintbrush and placing them back at the center, are inefficient, difficult to perform consistently, and too labor-intensive over long times, and thus not very practical. Longer experiments with adult *Drosophila* have successfully been automated to allow higher throughput (36–38), but no such system has previously been developed for freely crawling larvae.

Here, we present the design and operation of an automated “larva picker” robot and demonstrate its capabilities through continuous observation of larval exploration and navigation behavioral response with high detail and over unprecedented duration. Importantly, identity is maintained throughout tracking, so we can characterize exploration and navigation at the population and individual animal levels together, as larvae search for food under varying hunger conditions, or negotiate variable temperature environments. In doing so, we are able to reveal new behavioral dynamics, where the animals’ search strategy is modulated over hours and we can discriminate between different individual statistics that are otherwise hidden by population averages.

## Methods

### Larva Picker Robot

To perform long-term behavioral studies with the larva, we have designed and fabricated a robotic instrument that automatically tracks, transports, and feeds larvae throughout a large arena. The flexibility of its design enables arbitrary stimulus delivery alongside continuous measurement of behavior, yielding an unprecedented level of detailed locomotion data and a comprehensive view of locomotion strategies at the population and individual animal levels together.

The system operates by tightly coordinating video acquisition from an overhead camera with a manipulator arm capable of traversing three dimensions (Fig. 1B). The manipulator arm is translated by stepper motors (Nema 8) in the X- and Y-axes and a 5V solenoid (SparkFun) for the Z-axis, which are driven by a programmable controller board (SmoothieBoard) with physical and software limits in place to prevent overtravel. At the end of the arm is a custom 18-Gauge nozzle (Fig. 1C) that can interact with the larva and the experimental arena (Fig. 1D) that sit below the camera and manipulator assembly.

**Figure 1.**
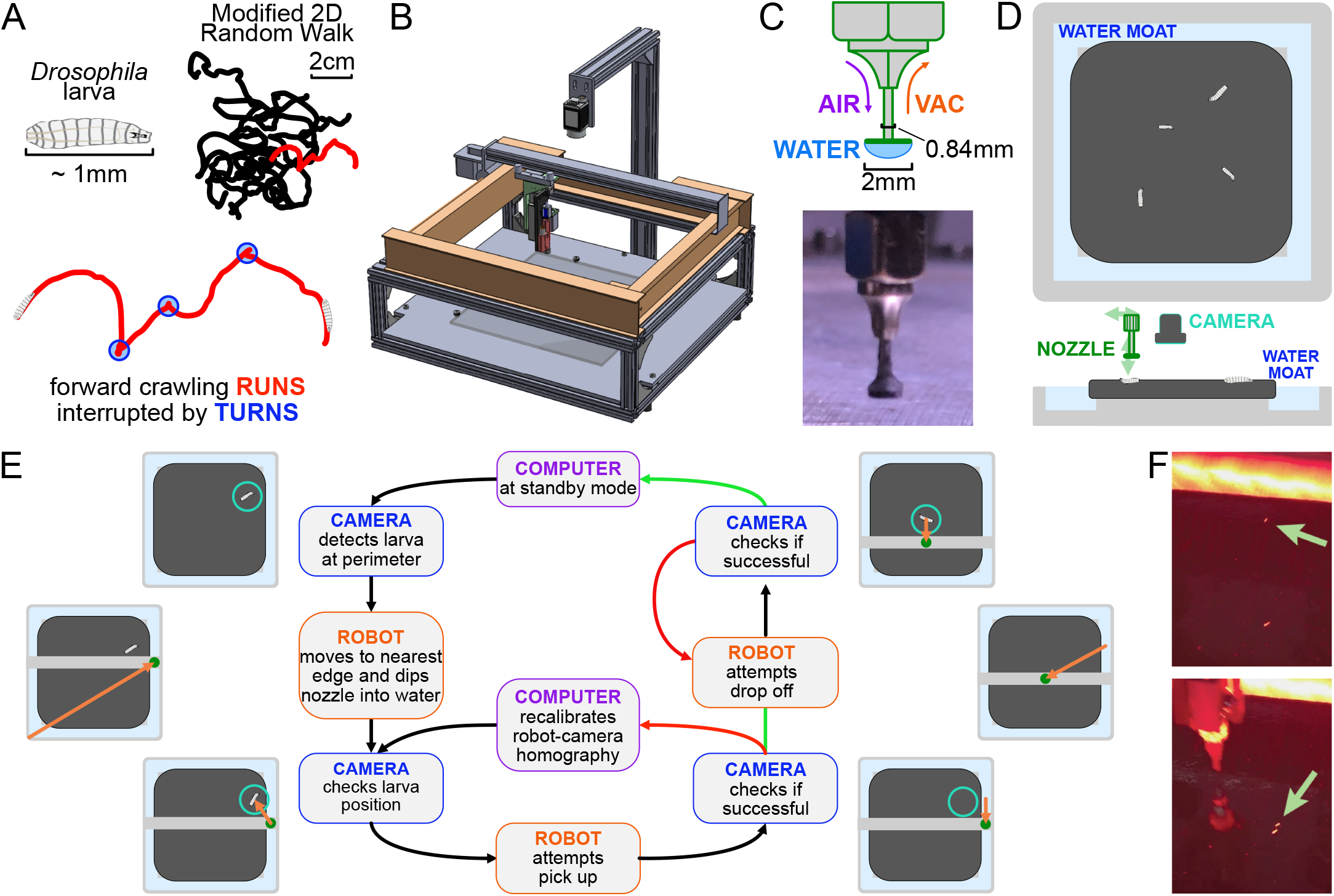
The automated larva transport robot enables continuous, long-term observation of fly larva crawling behavior. **A**. Schematic illustrating the fly larva’s simple search behavior. They explore their environment in a modified 2D random walk, with 20 example paths (black) shown. Trajectories are characterized by an alternating series of forward crawling runs (red) and turns (blue). **B**. Isometric CAD schematic of the transport robot. The robot is built from a modified 3D printer with a custom nozzle. Feedback from a mounted overhead camera allows for tight coordination with the moving arm to safely and robustly interact with the experimental arena. **C**. The nozzle is built as a narrow tube that allows air and vacuum flow with a flat plastic disk fitted at the bottom. The disk provides ample surface area for a water droplet to form, and the droplet’s surface pressure can pick up larvae while minimizing stress on the animal during the interaction. **D**. Top and side view schematics of the flat crawling arena. Larvae crawl atop an agar substrate, which is kept hydrated by a surrounding moat. The robot nozzle picks up larvae as they approach the edge of the arena and transports them back to the center to continue their free-roaming behavior. **E**. Flowchart of the larva pick-up feedback process. In standby mode, the camera records a video of larval behavior. When it detects a larva nearing the perimeter, it triggers the pick-up protocol for the robot. The manipulator arm moves to a point in the moat nearest to the larva and dips the nozzle in, forming a droplet at the tip to be used for pick up. The camera provides a more recent position for the moving larva as the robot attempts a pick up. If feedback from the camera suggests a successful pick up, it attempts a drop off. Otherwise, the manipulator repeats its attempt after receiving an updated larval position. Multiple failed attempts can trigger small perturbations to robot calibration parameters to allow better flexibility through reinforcement learning before continuing pick up attempts. Similarly, the robot performs multiple drop off attempts at the center of the arena until it receives a positive confirmation from the camera, at which point the system returns to its original standby mode. **F**. Photographs before (top) and after (bottom) the robot moves a larva from the perimeter to the center of the arena.

A 2.3-megapixel CMOS camera (Grasshopper3) observes a small number (4-6) of larvae crawling on a 22 × 22 cm agar gel (2.5% wt./vol. agar in water, with 0.75% charcoal added for improved contrast) and records at 10 Hz. Larvae are illuminated with four strips of red LEDs (dominant wavelength around 620 nm, which is outside the visible range of the larva (39), arranged in a square around the agar gel. To maintain larval exploratory behavior over a long duration, the camera detects when one nears the edge of the arena. This triggers the manipulator to pick up the larva with the nozzle. The larva is maintained on the nozzle via the surface tension of a water droplet. The water droplet provides a way to indirectly interact with the larva to prevent causing damage to the animal. After the manipulator arm moves to the center of the arena, it drops off the larva with a slow horizontal motion, effectively rolling it off the nozzle (Fig. 1E). The manipulator replenishes the water droplet before each pick up, and a small flat Delrin plastic disk (2 mm diameter) at the bottom of the nozzle provides more surface area for the droplet to form (Fig. 1C). When the surface tension of the water is insufficient to pick up the larva, the nozzle is capable of exerting vacuum suction to assist in pick up, as well as allowing air flow to release the larva during drop off.

Meanwhile, the mounted camera refreshes the image and therefore the position of the larva before and after each step of any protocol, ensuring that the process is robust to variations in crawling speed and behavior and to failed attempts, which are detected and repeated (Fig. 1E). The pick up/drop off procedure is 99.8%/99.9% successful (90%/95% on the first try), highly important for the viability of long-term experiments.

Our system must also address desiccation of the agar gel substrate and animal starvation, which limit experiment times and affect behavior. To address the former, we have built an auto-replenishing water moat (Fig. 1D) in direct contact with the gel, which then maintains its water content and physical shape. In addition, the moat also provides a convenient and readily available water source for the nozzle. To address the latter, the robot can automatically administer a drop of apple juice (≈ 0.1g/mL sugar concentration) directly to the larva on a predetermined schedule. The larva is allowed to eat for a fixed time, then rinsed with water so that it can return to a free-crawling state.

With these features in place, our system has so far achieved more than 30 hours of continuous observation of individual larva behavior.

In some experiments we observe how larvae navigate a variable sensory environment. We use a similar system as outlined in (6) to generate a 1D linear spatial thermal gradient. We fit the underside of the experimental arena’s aluminium base with hot and cold reservoirs on opposite sides, each equipped with two liquid-cooled water blocks. PID (proportional-integral-derivative) controllers drive thermoelectric coolers between each water block and its reservoir to maintain a thermal gradient of 0.035 °C/mm across the agar gel in the arena, with 13 °C on one side and 21 °C on the other side.

### Analysis Pipeline

Our data analysis pipeline extracts numerous behavioral features from video recordings of crawling larvae (Fig. 2). Custom computer vision software takes raw movies and extracts the position and body contour of each larva while preserving individual animal identities (Fig. 2A). Because the robot arm can briefly block the camera during pickup events, resulting in dropped frames, the software interpolates larval positions in these frames to maintain continuous observation. Since the larva’s body length is roughly 1 mm and it crawls with an average speed well below 1 mm/s, significant behavior events are unlikely to occur during dropped frames and interpolation is sufficient to rebuild trajectories.

**Figure 2.**
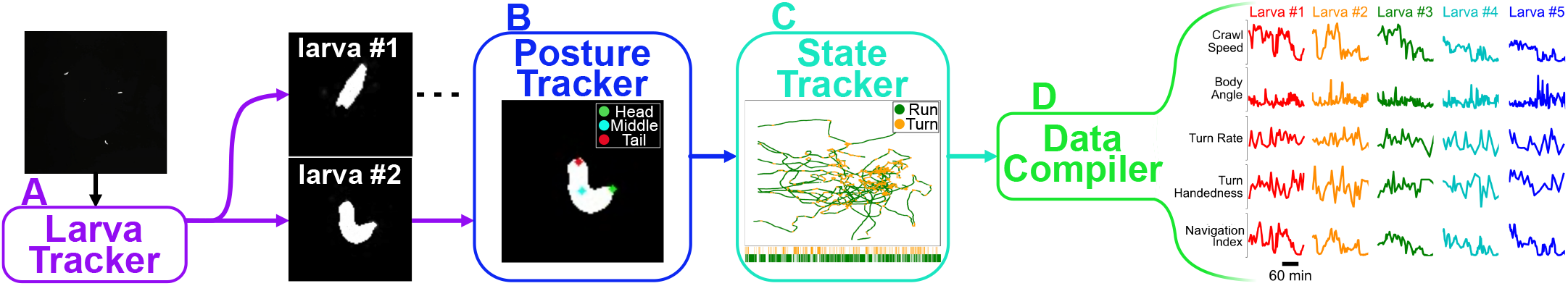
Analysis pipeline. **A**. Raw video acquired during the experiment is fed into computer vision software that tracks each larva while maintaining their identity. **B**. A posture tracker analyzes the isolated crops of each larva to determine its posture and orientation. **C**. A state tracker determines the behavioral state of each animal at each time point. **D**. Compiling all information from the preceding algorithms allows the pipeline to identify and calculate a wide variety of behavioral features.

The isolated crops (64 × 64 in pixels) of each larva are run through a recurrent U-Net (40) convolutional neural network (Fig. 2B) to determine the posture and orientation. The U-Net architecture has been shown to be highly effective at tasks that preserve spatial structure in an image by taking advantage of both global and local features (40–42). We design our model such that the output is a probability heat map of the head and tail of the larva, which preserves the global spatial structure of the larva’s body.

Traditional convolutional neural networks, including U-Net, often fail when analyzing sparse images with low-resolution features (41). This is particularly pronounced in our case since each larva is generally captured as clusters of only ≈ 30 uniformly bright pixels surrounded by black pixels. To compensate for this, we utilize more temporal information, such as the current momentum of the centroid. Since larval posture does not deviate much from frame to frame, we add recurrence in the form of long short term memory (LSTM) cells (43) at the beginning, middle, and final layers of the network to simplify the problem at each subsequent time step. The recurrence also creates an additional cost for head-tail flips which we have found to be a common issue with previous approaches to the problem (41, 44, 45).

Using all available information (position, contour, and posture), we use a densely connected neural network with bidirectional recurrence to classify the behavioral state of the larva (“run”, “turn”) at each time point (Fig. 2C) (34). The bidirectional recurrence here allows the network to identify the bends in the larva’s path and swings of the larva’s head by comparing frames both before and after. From here, we can identify a wide range of behavioral features and track them over extended time periods.

## Results

### The robot enables continuous observation of free exploration over six hours

Optimizing exploration by modulating behavior is essential to the larva’s ability to find a food source efficiently. While previous studies on *Drosophila* larvae have revealed some changes to its navigation over the first few minutes of exploration of an isotropic environment, how and whether their behavior evolves or persists afterwards remains unknown (7). Studies on another small organism with qualitatively comparable navigation dynamics, the nematode *C. elegans*, show similar changes in behavior during the first ≈ 100 seconds. Some longer-duration experiment (1 hour vs 15 minutes) reveal a transition in navigation strategy between two distinct modes of local and global searches, but transitions across similar or longer time scales have yet to be observed in *Drosophila* (7, 46). Here, we demonstrate how our larva picker robot enables continuous observation and analysis of larval locomotive behavior on very long time scales to study changes in its exploration strategy in an isotropic environment.

In Fig. 3, we present the results from continuous observation of second instar wild type (Canton-S) larvae (*N* = 42) freely roaming the experimental stage for a six hour time duration. Fig. 3A and C show the speed and the turn rate of the larvae, respectively, exhibiting the dynamics of their behavior. Notably, there is a significant drop in activity (both speed and turn rate) over the first hour before settling into a plateau that lasts for the following few hours. The correlation between larval crawl speed and turn rate over time has been previously observed (34) and is clearly evident here, and we measure a correlation coefficient of 0.572 in the population mean speeds and turn rates. With the large amount of data gathered on each individual, we confirm that this correlation also exist at the individual level with an average correlation coefficient of 0.348 ± 0.098.

**Figure 3.**
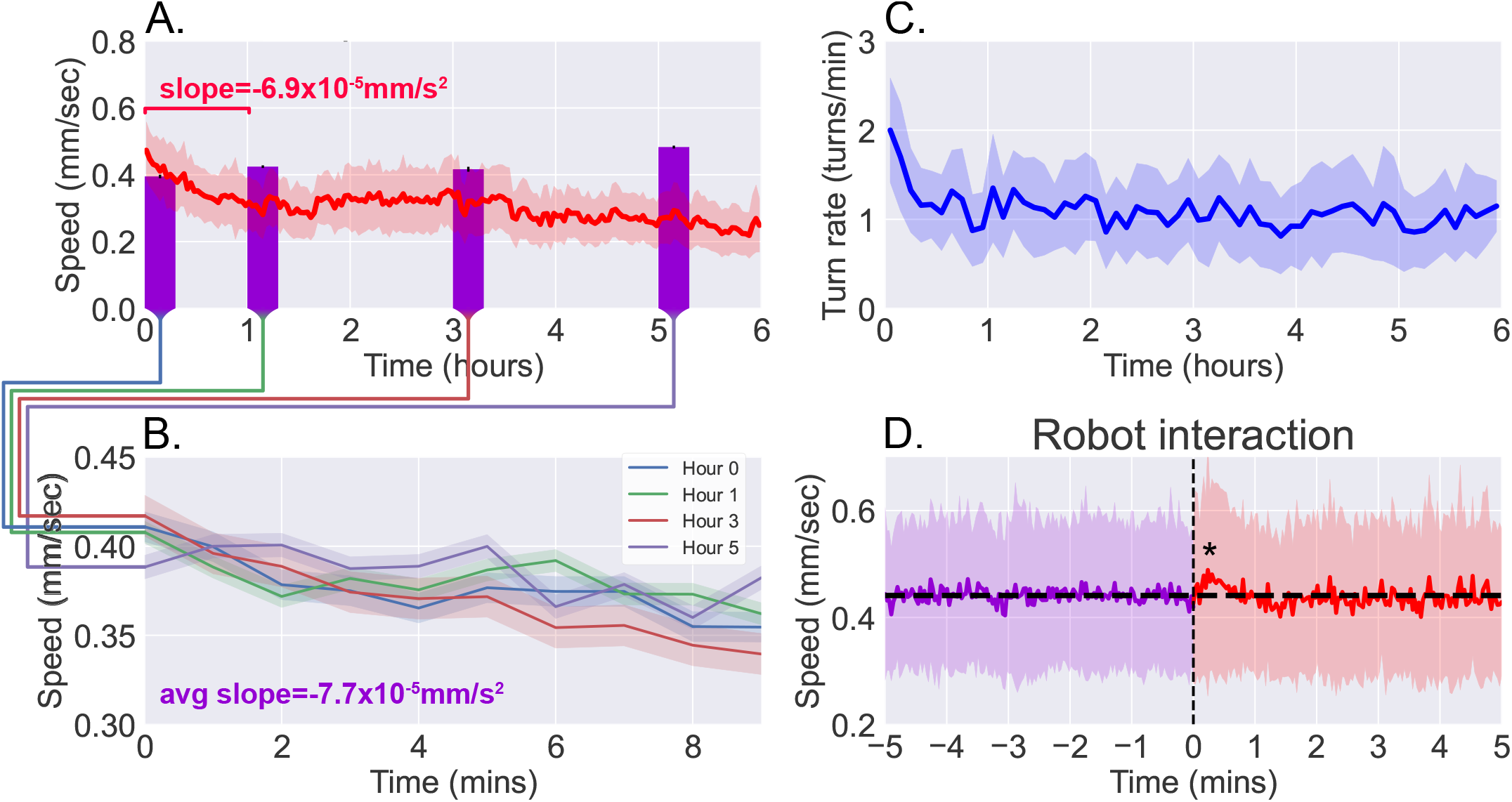
Observations of continuous free roaming for six hours. **A**. Larval crawl speed over time, comparing the results from continuous 6-hour observation recorded on the robot system (red, *N* = 42) to shorter 10-minute observation of larger numbers of starved larvae (purple, *N* = 200 per bar, from 10 experiments with 20 larvae each). **B**. Larval crawl speed over 10 minutes after starvation. We observe a decline in crawl speed (−7.7 × 10^*−*5^ mm/s^2^) comparable to that observed during the first hour of the continuous observation (−6.9 × 10^*−*5^ mm/s^2^). **C**. Larval turn rate over time. Similar to larval crawl speed (A), there is a noticeable drop in turn rate over the first hour, indicating an overall decrease in activity. **D**. Analysis of larval crawl speed before and after an interaction with the larva picker robot. We plot larval crawl speed during the five minutes immediately before (purple) and the five minutes immediately after (red) an interaction with the robot, i.e., a pick up and drop off event (vertical black dashed line). After a 1-2 min. transient (*p <* 0.05, Student’s t-test), speed returns to the mean pre-interaction level (horizontal black line). For all panels, shaded regions indicate standard deviation from the mean.

### Long-term free roaming behavior is consistent with analogous snapshot observations

Since the larva were not fed over the duration of the experiment, we compare continuous observations with short “snapshot” observations of larvae at various stages of starvation (*N* = 200 per stage). Larvae were removed from food and starved over a certain number of hours, then placed on an agar gel arena to be observe for 10 minutes without interruption, and with no interaction with the robot. The decline in activity seen in the robot-mediated experiments is not present when observing crawl speeds averaged over the 10-minute duration (Fig. 3A, purple bars). Upon closer inspection, however, larval crawl speed measured in the starvation studies (Fig. 3B) reveal a trend over time that mirrors what we observe in the first hour of continuous observations. Over these 10-minute snapshot observations, we measure an average decline of −7.7 × 10^−5^ mm/s^2^, comparable to the average of −6.9 × 10^−5^ mm/s^2^ seen over the first hour of continuous observations. The similarity offers confidence that the behavioral dynamics present here are not caused by transport robot actions but are real features of the animal that that continue to persist and develop over a period that is more than 30 times longer than what snapshot observations can capture.

To further ensure that disruptions to behavior due to the robot’s interference have minimal influence, we analyzed larval behavior before and after each interaction with the robot (i.e., a pick up and drop off event). Fig. 3D shows averaged larval crawl speeds 5 minutes before and 5 minutes after an interaction event (vertical dashed line). As expected, mean larval speeds before the interaction shows little change over the small window. We do observe a transient of approximately 1-2 minutes just after the interaction, where there is a noticeable increase in speed (p<0.05), but it quickly returns to the same mean as before (horizontal line).

### A freely-crawling single larva is continuously monitored for more than 30 hours

As noted in the previous section, the robot is also capable of automatic scheduled feeding. To keep the larva alive as long as possible, the robot administers a drop of sugar-rich solution directly to the larva once every hour. The larva is allowed to feed for one minute, then the animal and the drop site is rinsed with water, and the larva resumes exploration. Using this protocol, we are able to study larval locomotive behavior for over 30 hours, yielding an unprecedented amount of behavioral and developmental information on an individual animal. Fig. 4 presents the trajectory of a single larva and some of its behavior features observed over a duration of 30 hours, where the individual animal crawls for more than 48 meters. Some behavioral features exhibit steady change, like a decreasing speed throughout the experiment. Notable differences in path shape occur approximately half way through the long observation, with dramatic increases in curvature and turn size, consistent with the tight loops seen in the full trajectory plot (Fig. 4 left). We also note that this larva maintains a left-turning bias throughout the experiment.

**Figure 4.**
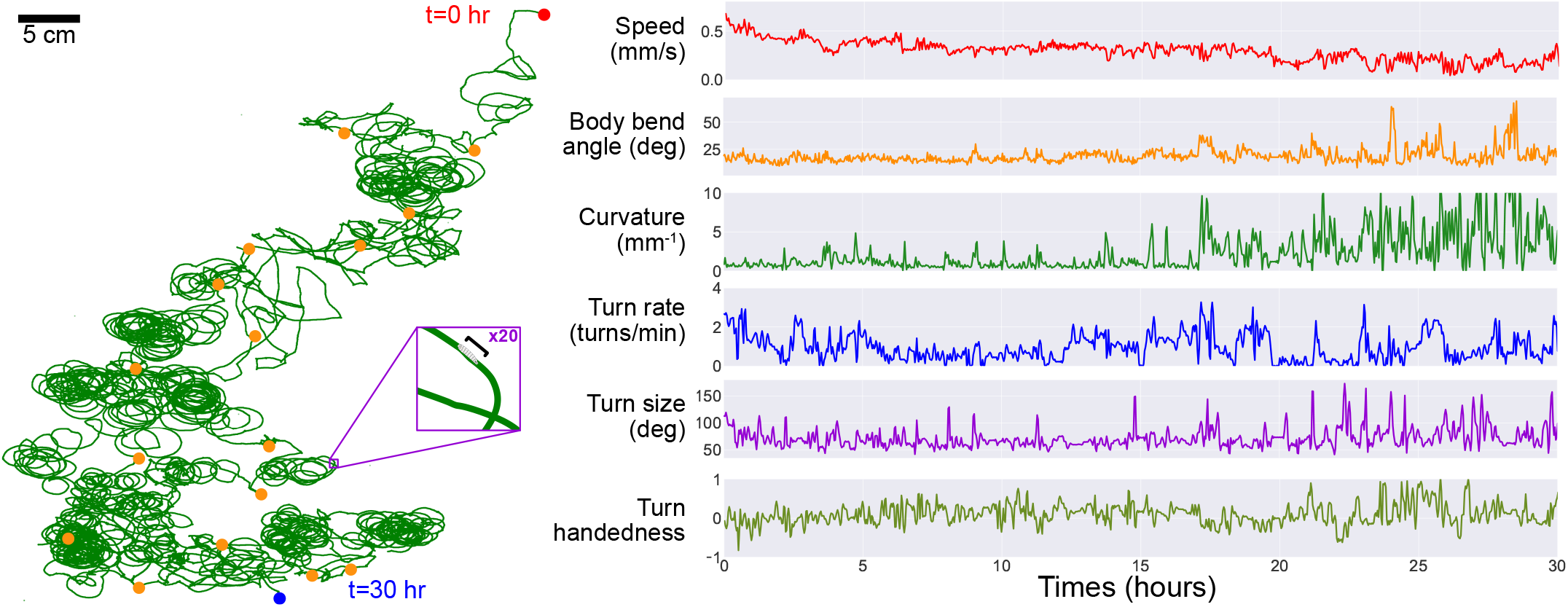
Long-term observation of a single larva. In order to maintain exploratory behavior in a fly larva without starving it, the robot automatically delivers a drop of apple juice (≈ 0.1g/mL sugar concentration). The larva is allowed to eat for 1 − 2 minutes, after which the robot uses water and air to rinse the larva, which then continues roaming freely. This protocol allows for continuous observation of larval behavior over developmental timescales. **Left**. The larva’s trajectory over a 30-hour duration, with its path (green) stitched together at each robot pick-up and drop-off positions (orange markers) to produce a continuous trajectory. The larva begins at the top right (red) and ends its run at the bottom (blue). **Cutout**. A x20 magnification on a small section of the path, showing the scale of the path compared to the larva’s body (black bar, ≈ 1mm). **Right**. We plot a number of behavioral features observed during its exploration, including its speed (red), body bend angle (orange), trajectory curvature (green), turn rate (blue), turn size (purple) and turn handedness (olive). Turn handedness is calculated as (*N*_*left*_ − *N*_*right*_)*/N*_*total*_, such that a handedness of 1.0 indicates all left turns, and a handedness of −1.0 indicates all right turns.

### Larval thermotaxis is maintained over long periods

By leveraging the automated transport system’s flexible design to deliver and study responses to stimuli, we manipulate the temperature of the agar arena to study larval thermotaxis behav-ior. *Drosophila* larvae have robust, highly-sensitive, and well-documented response to changes in temperature, and work has been done to decipher the behavioral strategy utilized to efficiently navigate thermal gradients (6, 24). However, there is a lack of abundant data on individual animals, since single experiments last on the order of 10 minutes, and thus a lack of understanding of the differences in thermotaxis strategy and its development between individuals. It is also unknown how thermotaxis might evolve over long times. The automated transport system here provides an opportunity to delve deeper into these questions.

Navigational effectiveness can be captured by a dimensionless navigation index equal to ⟨*v*_*x*_⟩ */*⟨*v*⟩, the average of the component of crawling velocity along the thermal gradient normalized to the average speed. The resulting thermal navigation index ranges from +1.0 (parallel to gradient, i.e. crawling directly towards the warm side of the arena) to – 1.0 (anti-parallel to gradient, i.e. crawling directly towards the cold side of the arena) (6, 24, 47). Fig. 5 shows the thermal navigation index of second instar wild type (Canton-S) larvae as they crawl across the experiment arena for six hours, both with (*N* = 38) and without (*N* = 42) a thermal gradient present. The thermal gradient is centered at 17 °C with a steepness of 0.032 °C/mm, which would normally evoke robust cold avoidance behavior (24).

**Figure 5.**
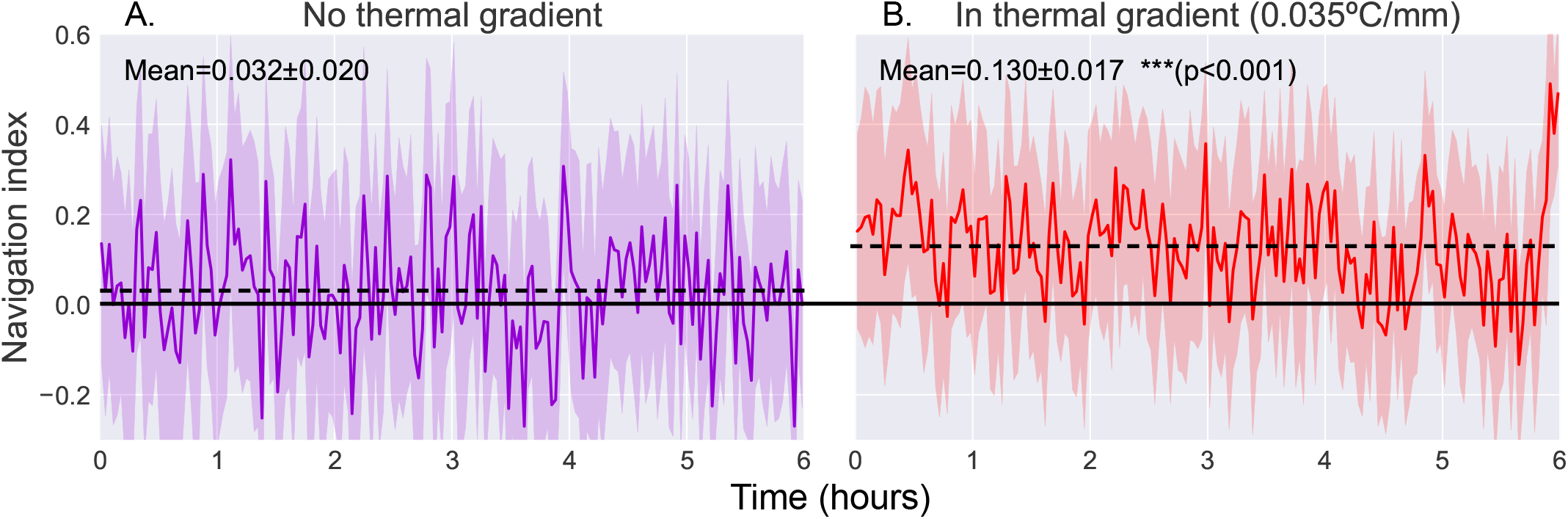
Comparison of navigation index in a zero gradient environment (**A**, *N* = 42) and in a presence of a linear thermal gradient of 0.035°*C/mm* (**B**, *N* = 38). Thermal navigation index is calculated as a dimensionless index equal to ⟨*v*_*x*_⟩*/*⟨*v*⟩, such that +1.0 is parallel to gradient, 0.0 is normal to gradient, and −1.0 is anti-parallel to gradient. We observe a clear increase in average navigation index when exposed to a thermal gradient (increase from 0.032 ± 0.020 to 0.130 ± 0.017 (*p <* 0.001, Student’s t-test)), but we do not observe any significant pattern of change in that index over time. Shaded region indicates one standard deviation.

When roaming freely with no stimulus present, we observe an average navigation index of 0.032 ± 0.020 (range indicating standard deviation). When a thermal gradient is applied to the arena, we observe a clear increase in navigation index to an average of 0.130 ± 0.017 (*p <* 0.001), indicating positive thermotaxis, where larvae crawl towards warmer temperature. The navigation index also remains steady over the 6-hour measurement time.

### Navigation efficiency of individuals exhibits distinct behavioral phenotypes

Short snapshot observations produce limited information on any single individual animal, therefore limiting most thermotaxis analysis to population-level statistics. Long continuous observation enabled by the transport robot produces much more detail on individual animals, allowing us to analyze behavioral features at the individual level as well (Fig. 6). Importantly, averaging over a population necessarily results in some loss of information such that different statistics at the individual level can generate the same population mean.

**Figure 6.**
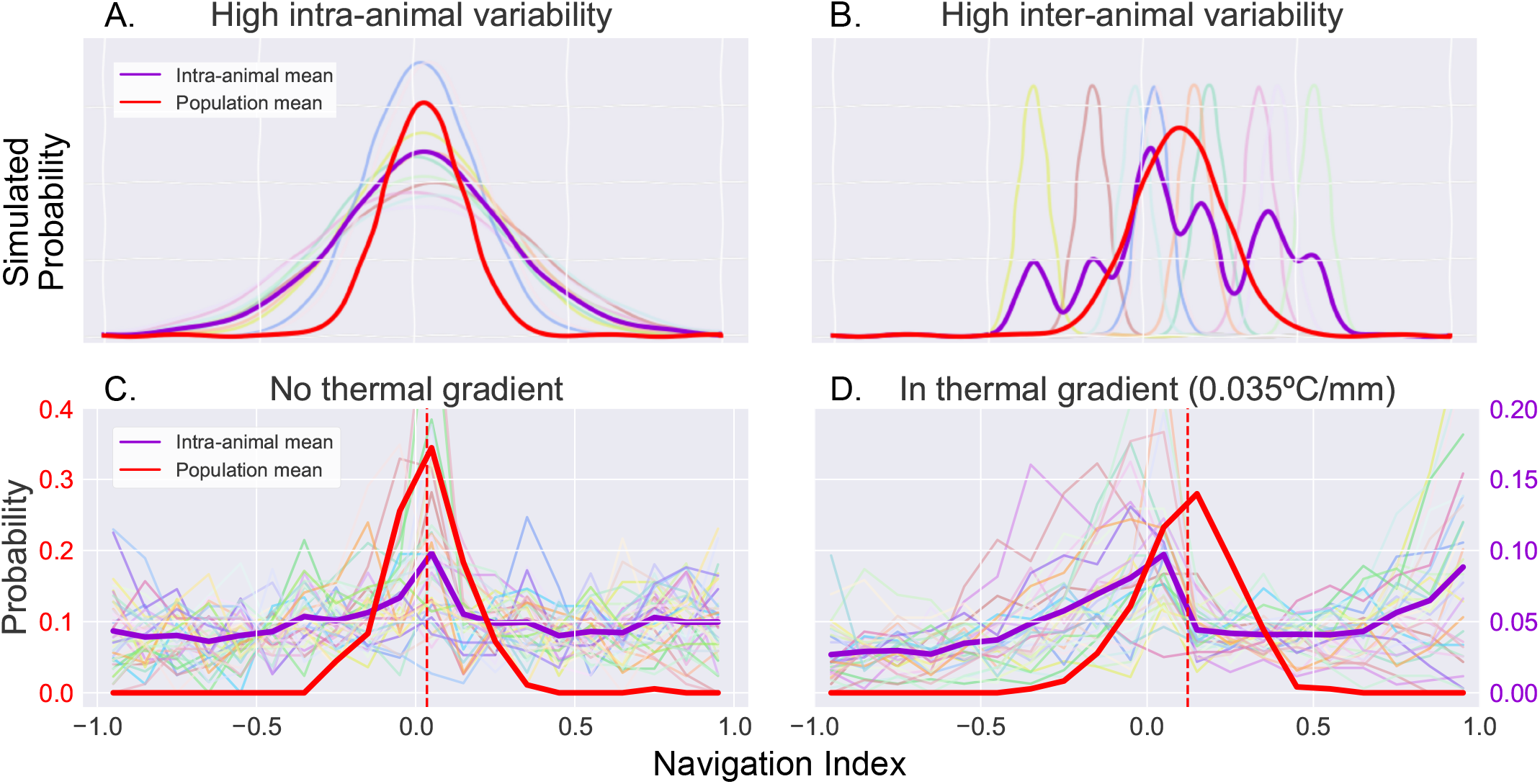
Examination of inter- and intra-animal variability via analysis of probability distribution of the observed larval thermal navigation index. When dissecting probability distributions of observed behavior, we notice that the same population (inter-animal) mean can be produced by two individual (intra-animal) distributions. **A**. Simulated example of individual probability distributions with high intra-animal variability. The resulting mean of intra-animal distribution (purple) closely resembles the population mean (red). **B**. Simulated example of individual probability distributions with high inter-animal variability. The resulting mean of intra-animal distribution (purple) form a multimodal distribution despite a similar population mean (red) as A. **C**. Probability distribution of thermal navigation index observed without a thermal gradient. There is high intra-animal variability but low inter-animal variability, such that the intra-animal mean forms a similar distribution to the population mean, as was the case seen in A. **D**. Probability distribution of thermal navigation index observed in presence of a thermal gradient (0.035° C/mm). In contrast to C, the intra-animal mean forms a bimodal distribution, more closely resembling a distribution with high inter-animal variability as seen in B.

Fig. 6A-B demonstrates the differences between two such cases by examining simulated toy examples of a probability distribution of observations of a navigation index. Each series of observations is sampled from a Gaussian distribution, whose mean and standard deviation are randomly determined with two different statistics. Both simulations have similar distributions when analyzing navigation index occurrences at the population level (red), but produce distinct means of intra-animal probability distributions (purple). In the first simulation (Fig. 6A), high intra-animal variability produces a mean of intra-animal distributions that is similar to the population mean distribution. In the second (Fig. 6B), high inter-animal variability instead pro-duces a multimodal distribution for individuals despite the extra modes not being present at the population level. Because our long time scale thermotaxis experiments produce enough data to establish both population-level and invididual-level navigation indexes, we can examine thermal navigation in a similar way.

Interestingly, we find that larval thermotaxis behavior switches between the two models when a thermal gradient is applied (Fig. 6C,D). While the population means have a similar shape and spread in both contexts (red traces), without a thermal gradient (Fig. 6C), the individual distributions exhibit high intra-animal variability and low inter-animal variability, mirroring those seen in the former (A). When in the presence of a thermal gradient (Fig. 6D), we observe a bimodal distribution that more closely resembles the latter (B) instead. We measure a binomial coefficient (BC) (33, 48) of 0.67 here, compared to *BC* = 0.48 when there is no gradient present. This is a significant increase (*p <* 0.01) that crosses the critical value for detecting bimodality (*BC*_*crit*_ = 5*/*9), clearly indicating a shift in the shape of the distribution.

## Discussion

We have developed an automated system for long-term observation of *Drosophila* larvae (Fig. 1). The robotic arm is capable of transporting larvae as they approach the edge of the experiment arena back to the center, allowing continuous observation of exploratory or directed navigation behavior from an overhead camera. Through coordination and constant feedback between the robot and video acquisition, the system maintains larvae within the arena with high reliability, and we are able to achieve continuous observation over developmental timescales. The accompanying analysis pipeline takes the output video from these experiments to track larval posture and behavioral state while maintaining individual identities. The analysis compensates for the output video’s low resolution and lack of detailed features of the larvae through a combination of local and global features and the use of recurrent neural networks (Fig. 2).

We present a study of free roaming behavior in larvae over six hours of continuous observation, comparing results to short 10-minute snapshot observations of larvae at various stages of starvation (Fig. 3). The comparison yields dissimilar patterns when considering the 10-minute averages, but similar dynamics when considering the change in behavior over time, such that the 10-minute trajectory in larval crawl speed resembles that seen in the first hour of continuous observation. The similarity suggests behavioral dynamics that are present in both experiments, but persist and develop over a duration that is an order of magnitude longer than what snapshot observations can capture.

Since our analysis maintains animal identities throughout the video, we are able to capture behavioral information on single individuals with unprecedented detail. We leverage this new trove of data to analyze larval response to a thermal gradient and examine the probability distribution of the observed navigation index over time (Fig. 6). In particular, we note that the same population (inter-animal) mean can be produced by different individual (intra-animal) distributions. Interestingly, larvae seem to switch between two such distribution shapes upon encountering a thermal gradient. The individual distributions of thermal navigation index without a thermal gradient exhibits high intra-animal variability but low inter-animal variability, forming a unimodal mean distribution that is similar to the population mean. In contrast, the individual distributions exhibits high inter-animal variability, such that it become bimodal when a gradient is applied, despite the distribution shape remaining unimodal at the population level. This is consistent with recent findings that suggest a switch-like (all-or-none) learning behavior in larval *Drosophila*, which is also not apparent when only analyzing population means (49). By observing decision-making behavior of individual larvae over several cycles of stimulus and reward, *Lesar, et al*. (49) find that Pavlovian training of preference for carbon dioxide is similarly quantized to two states (all-or-none), each centered at a fixed preference index.

While the context and modality for the learning assays are different than our observation of larval thermotaxis, both results reveal new features of larval behavior that were previously obscured in population averages. Both results also suggest the existence of larval “personality types”, or distinct, quantized behavioral phenotypes that vary between individual animals (6, 34, 49, 50). This work provides some progress in uncovering such phenotypes and furthering our understanding of learning and development of navigational strategies in individual animals in addition to population averages.

With the robot’s flexible design, we can continue to probe these questions in many different contexts. For example, the robot can deliver food or soluble drugs directly to the larvae on a predetermined schedule to measure both the acute and chronic effects on the animal’s behavior and physiology (27, 51–56). We can also leverage the existing lighting system to implement optogenetic activation or suppression of specific neurons, allowing delivery of fictive stimuli or studies of the effects of certain neuronal circuits on larval behavior and development (4, 5, 19, 57, 58). This can either be activated on a predetermined schedule, or integrated directly with the existing robot and camera feedback system to enable activation triggers based on specific conditions such as larval behavior (e.g. activate upon larva initiating a turn).

A major obstacle in way of achieving continuous observation of the entire larval life cycle (≈ 100 hours) is a reliable method of delivering enough nutrition and ensuring the larva has sufficiently fed. The current method of delivering drops of sugar-rich solutions has not been sufficient to trigger molting into the next instar stage, thus limiting us to a single instar stage. We hope to solve this problem and others as we continue to develop the system and expand its capabilities, for example with a second adjacent arena with more nutritive food where larvae could experience longer feeding times before being returned by the robot to the primary behavioral arena. Such rich detail covering the entire development of the larva would provide powerful insight into the long-term learning, memory, and behavioral adaptations in tandem with the physiological developments that take place between instar stages.

We hope that this study has highlighted some advantages of long term continuous measurement, and that similar instrumentation and analysis method could be applied to other organisms, along with a wide range of investiations in the fly larva.

## Acknowledgements

VV acknowledges support from the Burroughs Wellcome Fund and NIH R01 NS126334. MK acknowledges support from the NSF CAREER Award 2144385.

## Data Availability

Code written for this paper can be found on GitHub here https://github.com/venkatachalamlab/LarvaPickerRobot.. Mildly compressed versions of all video recordings can be found hosted on Zenodo.

